# Exploring Gene-Mediated Mechanisms Behind Shared Phenotypes Across Diverse Diseases Using the clGENE Tool

**DOI:** 10.1101/2024.04.08.588642

**Authors:** Li Zheng

**Affiliations:** Zhongshan Institute for Drug Discovery, Shanghai Institute of Materia Medica, Chinese Academy of Sciences, Zhongshan 528400, P. R. China

**Keywords:** Shared Molecular Mechanisms, Multi-Omics Data Integration, Phenotype Analysis, Data Integration

## Abstract

The observation of similar clinical characteristics across a broad spectrum of diseases suggests the existence of underlying shared molecular mechanisms. Identifying these mechanisms is critical for uncovering the molecular roots of diseases and advancing the development of innovative therapeutic strategies. However, researching the common genes that mediate similar phenotypes among different diseases often requires the integration of various sequencing datasets and clinical data. The batch effects among these datasets and the complexity of clinical data present significant challenges to the research. This study developed a framework named “clGENE”, aimed at uncovering the molecular mechanisms behind similar phenotypes across different diseases. By integrating data normalization, cosine similarity analysis, and principal component analysis (PCA) algorithms, this framework is capable of effectively identifying shared molecular mechanisms associated with specific phenotypes and further selecting key shared genes. Through the analysis of a pan-cancer dataset, we have verified the efficacy and reliability of the “clGENE” framework. Furthermore, this study also established a dataset on immune cell infiltration and successfully identified key patterns of immune cell infiltration in different cancer lymph node metastasis stages using the ‘clGENE’ framework, further confirming its potential application in biomedical research.

## Introduction

Present research has illuminated that numerous diseases exhibit significant phenotypic similarities[1, 2]. This suggests the possibility of shared molecular pathways and gene-mediated mechanisms across different diseases. Uncovering these shared molecular mechanisms often necessitates the integration of multiple sequencing datasets[3, 4]. With the revolutionary advances in high-throughput technologies, the cost of generating omics data has significantly decreased, which has facilitated the emergence of numerous public databases collecting various types of sequencing data[5, 6]. This shift has opened new avenues for exploring potential shared molecular mechanisms across different diseases.

Unveiling the shared molecular mechanisms across various diseases is essential for developing therapeutic strategies applicable to multiple conditions. However, this endeavor faces notable challenges related to technology and data processing methods. Among these, the batch effect issue, arising from processing biomedical data from diverse sources, stands out as a considerable obstacle[7]. The inherent variability in biological samples, disease states, experimental conditions, and sequencing technologies leads to data that can significantly differ in magnitude, precision, and scope[8]. Such variability may not only mask genuine biological signals but may also lead to erroneous interpretations, thereby hampering the identification and analysis of shared genes or biological pathways[9]. Additionally, in the realm of sequencing data and clinical information, phenotypic details are frequently entangled[10]. For instance, a disease’s clinical presentation might encompass both dermatological symptoms and immune system irregularities, with these phenotypic elements being interwoven. This complexity underscores the importance of precisely pinpointing the specific molecular mechanisms tied to particular phenotypes, whether they be gene mutations associated with dermatological manifestations or molecular pathways linked to immune system disturbances.

Addressing this challenge, the present study developed clGENE, an advanced computational framework aimed at identifying and analyzing genes shared across different diseases associated with specific phenotypes. By integrating data normalization, cosine similarity analysis, and Principal Component Analysis (PCA), clGENE successfully navigates batch effects, accurately pinpointing shared molecular mechanisms and key molecular markers across diseases. Furthermore, the study validates the application potential of clGENE using a pan-cancer dataset and demonstrates its efficacy in recognizing patterns of immune cell infiltration at various cancer stages through the construction of an immune cell infiltration dataset. The R package is available for download at https://github.com/lizheng199729/clGENE.

## 2. Materials and Methods

### 2.1 clGENE Algorithm

Initially, each feature in the raw dataset is subjected to standardization, ensuring that the standard deviation of each feature is set to 1. (Figure 1B)

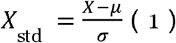

**Figure 1:**
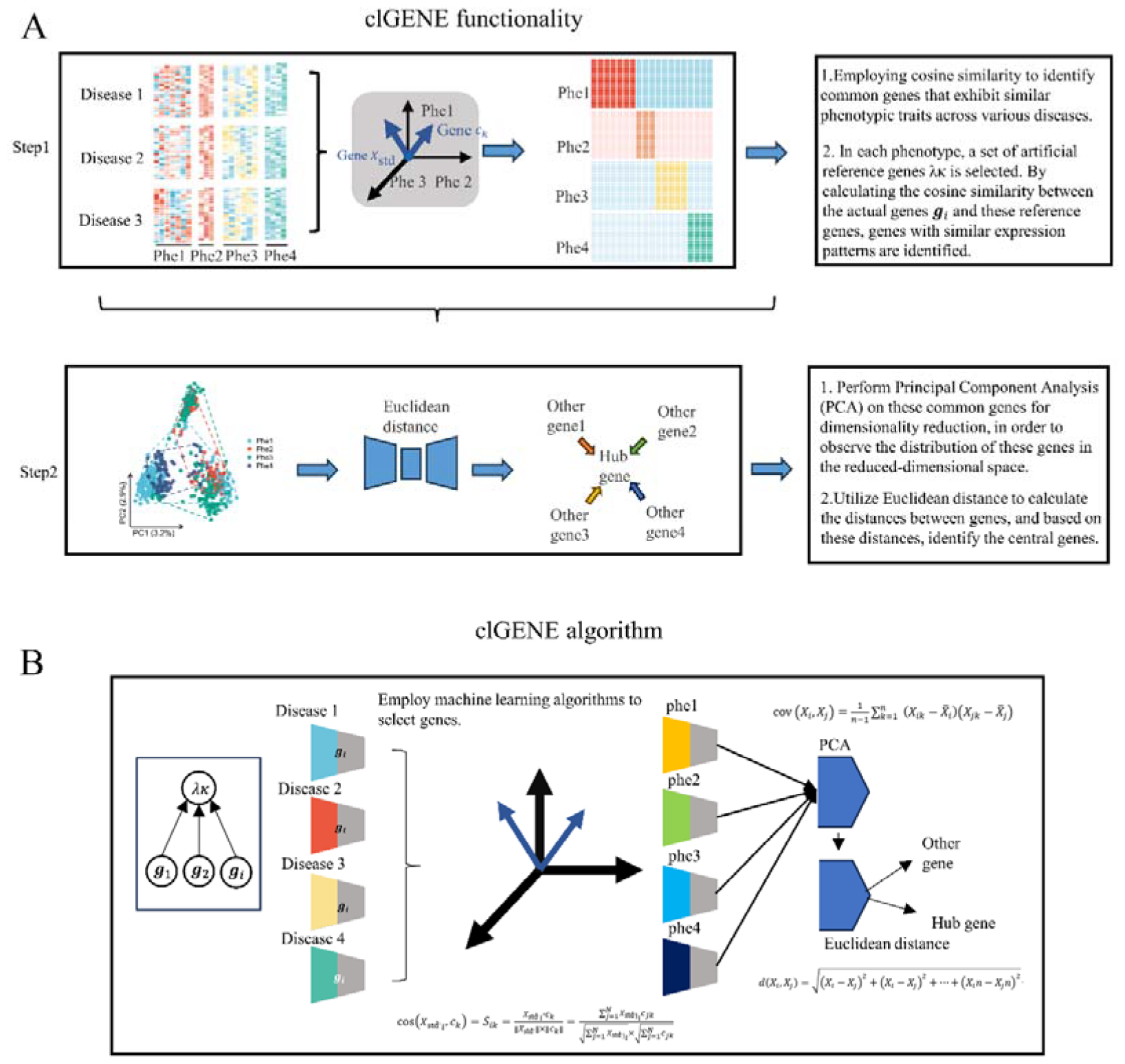
Overview of the clGENE algorithm. A. Framework and functionality of clGENE. B clGENE operates under the hypothesis that there exists a common molecular mechanism underlying similar phenotypes across different diseases, a notion difficult to unveil due to the high-dimensionality of omics data and the complexity of clinical data. Initially, clGENE employs L1 regularization for data preprocessing. Subsequently, leveraging clinical data for phenotype grouping, it employs a cosine similarity algorithm to identify potential molecular mechanisms mediating common phenotypes. Further, clGENE applies PCA dimensionality reduction to each potential molecular mechanism, and utilizes Euclidean distance for gene clustering, thereby filtering for gene clusters associated with potential molecular mechanisms, aiming to reveal the molecular basis behind similar phenotypes across varying diseases.

Where *X* represents gene expression data, *µ* is the mean of the gene expression data, *σ* is the standard deviation of the gene expression data, and *X*_std_ is the standardized data.

In the subsequent analysis, we employ L1 regularization techniques to conduct an in-depth investigation of the standardized gene expression matrix. This step is a common practice in statistical modeling, particularly suited for handling high-dimensional datasets, such as gene expression data, as denoted by the formula.

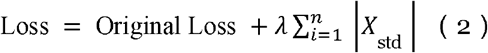

The model’s loss function, designed to be minimized, combines the original loss with an L1 regularization term. The original loss assesses prediction accuracy, while the L1 term, weighted by a regularization parameter λ, adds a penalty for the magnitude of the model’s coefficients, promoting sparsity. Specifically, for a gene expression matrix, the L1 regularization term is the sum of the absolute values of the standardized coefficients, thereby aiding in feature selection within high-dimensional datasets.

Subsequently, to screen for potential genes associated with the same phenotype across different diseases, we employed a cosine similarity algorithm. Specifically, we introduced an artificial gene that is expressed as 1 only in samples with the same phenotype, while it is expressed as 0 in all other samples.

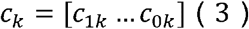

The next step involves determining the cosine similarity between 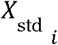 and *c*_*k*_.

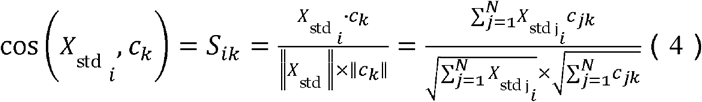

Subsequently, to more thoroughly quantify and comprehend the linear relationships within gene expression data, we will calculate their covariance matrix.

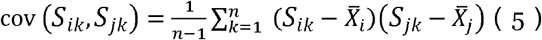

*S*_*ik*_ represents the gene expression discovered through cosine similarity, and *S*_*jk*_ is the expression of other genes. *X*_*i*_ and *X*_*j*_ are the sample means of variables *S*_*ik*_ and *S*_*jk*_ respectively. *n* denotes the number of samples.

We utilize the Euclidean distance to calculate the proximity between the central gene and other genes, thereby identifying key genes closely associated with specific biological states or disease stages. The Euclidean distance, a common metric for measuring the straight-line distance between two points, can reflect the degree of similarity or dissimilarity in gene expression patterns. By analyzing the Euclidean distances between genes, we are able to effectively unearth those that have significant associations with particular biological processes or pathological states. The specific expression is as follows.

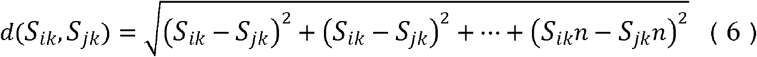

### 2.2 Gene enrichment analysis

KEGG pathway enrichment analysis was performed with clusterProfiler with a background set of all entrez ids mapped to a KEGG pathway.

### 2.3 Pan-cancer dataset

The pan-cancer dataset, sourced from The Cancer Genome Atlas (TCGA), compiles clinical and gene expression data from 2,452 cases across various cancer types, including Adrenocortical Carcinoma (ACC), Bladder Urothelial Carcinoma (BLCA), Breast Invasive Carcinoma (BRCA), Glioblastoma Multiforme (GBM), and Kidney Renal Papillary Cell Carcinoma (KIRP).

### 2.4 Analysis of immune cell infiltration

This study utilizes ssGSEA and CIBERSORT, methodologies previously elucidated in published articles[11].

## Result

### clGENE introduce

clGENE represents an advanced machine learning model that, by analyzing multiple dimensions including transcriptomics and proteomics, uncovers the shared mechanisms behind similar phenotypes across diverse diseases. This model is predicated on the foundational hypothesis that there exists a similarity in phenotypic mechanisms across various diseases, thereby suggesting the presence of shared molecular mechanisms. Furthermore, this hypothesis encompasses the notion that there is a certain degree of interconnectivity among genes, which plays a pivotal role in the construction and functional expression of biological networks. In this model, the analytical process is distinctly divided into two primary phases.

Initially, by employing the cosine similarity algorithm, we are capable of identifying specific genes with unique expression patterns in diseases that exhibit similar phenotypes. Given the significant variability and complexity often inherent in sequencing data, particularly when it integrates clinical information from a multitude of diseases, the cosine similarity algorithm emerges as an effective instrument for meticulously managing and interpreting this complexity. The second part of the analysis involves conducting Principal Component Analysis (PCA) for dimensionality reduction, coupled with the use of Euclidean distance for the identification of key gene clusters. This approach is designed to streamline the complexity inherent in gene expression data by reducing its dimensions while preserving essential variability information, thereby enhancing the interpretability of the data and elucidating the relationships between genes. This process not only aids in pinpointing genes and gene clusters that play pivotal roles in disease progression and phenotypic expression (Figure 1A).

clGENE is an R package developed for the analysis of various omics data, facilitating the integrated analysis of multi-omics datasets. Its primary objective is to assist researchers in uncovering the potential molecular mechanisms underlying similar phenotypes across different diseases (Figures 2A and 2B). clGENE requires the input to be a standardized expression matrix of genes, proteins, or metabolites, with the first row designated as “group” for clinical phenotype data (Figure 2C). Utilizing this clinical phenotype information, clGENE automatically identifies gene expressions specific to particular clinical phenotypes and employs PCA algorithms along with Euclidean distance measures to filter gene clusters associated with potential molecular mechanisms (Figure 2D). Moreover, clGENE offers three visualization methods and provides access to its source code, further supporting researchers in their analysis and investigative efforts (Figure 2E).

**Figure 2:**
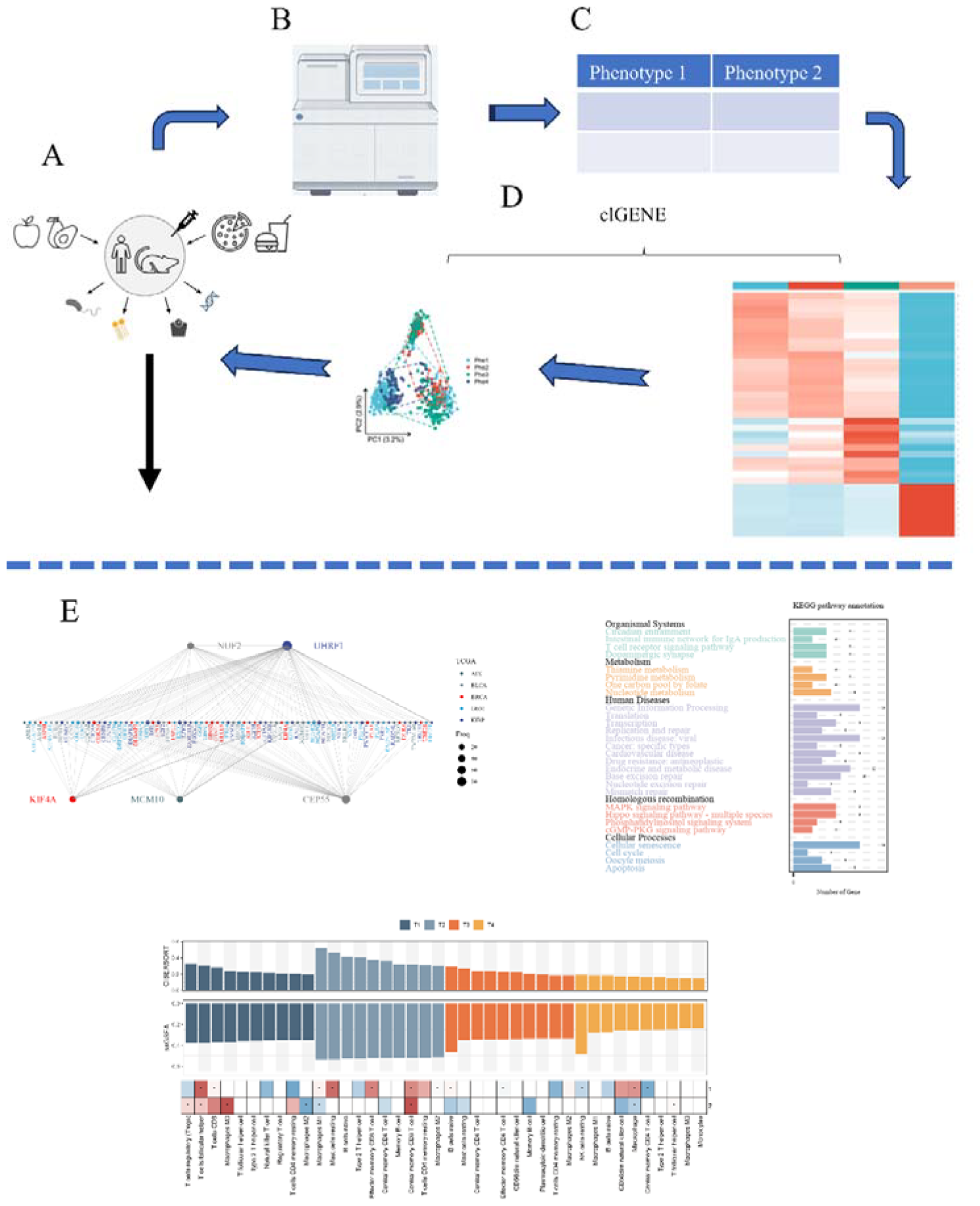
delineates the application process of clGENE. A,B depict omics data from various diseases in humans or animals. C indicates that clGENE requires the input of omics data expression profiles along with clinical phenotype information from different diseases. D elaborates that clGENE assists users in identifying key genes and gene clusters associated with common phenotypes across different diseases. E, clGENE offers three distinct forms of visualization to facilitate a deeper understanding and interpretation of the connections between these key genes and gene clusters.

### Identifying Key Expression Genes of Common Phenotypes Across Different Diseases Through clGENE

To assess clGENE’s effectiveness in pinpointing crucial gene expressions, our research incorporated a publicly accessible pan-cancer dataset. This dataset was segmented into lymph node metastasis based on clinical data. The application of Uniform Manifold Approximation and Projection (UMAP) for visual analysis showcased clGENE’s proficiency in managing intricate datasets spanning multiple diseases. Initial UMAP findings indicated a blend of gene expressions across various stages (as depicted in Figure 3A, left side). Contrasting this, post-clGENE application results highlighted a pronounced delineation in key gene expression traits specific to each stage (shown in Figure 3A, right side). Additionally, heatmaps revealed strong correlations between clGENE-filtered genes and clinical phenotypes, thus efficiently pinpointing genes linked to particular phenotypes (Figure 3B). Further, box plots detailed the expression dynamics of these identified key genes across differing phenotypes, solidifying clGENE’s effectiveness in isolating genes intimately connected with specific phenotypes (Figures 3C-F).

**Figure 3:**
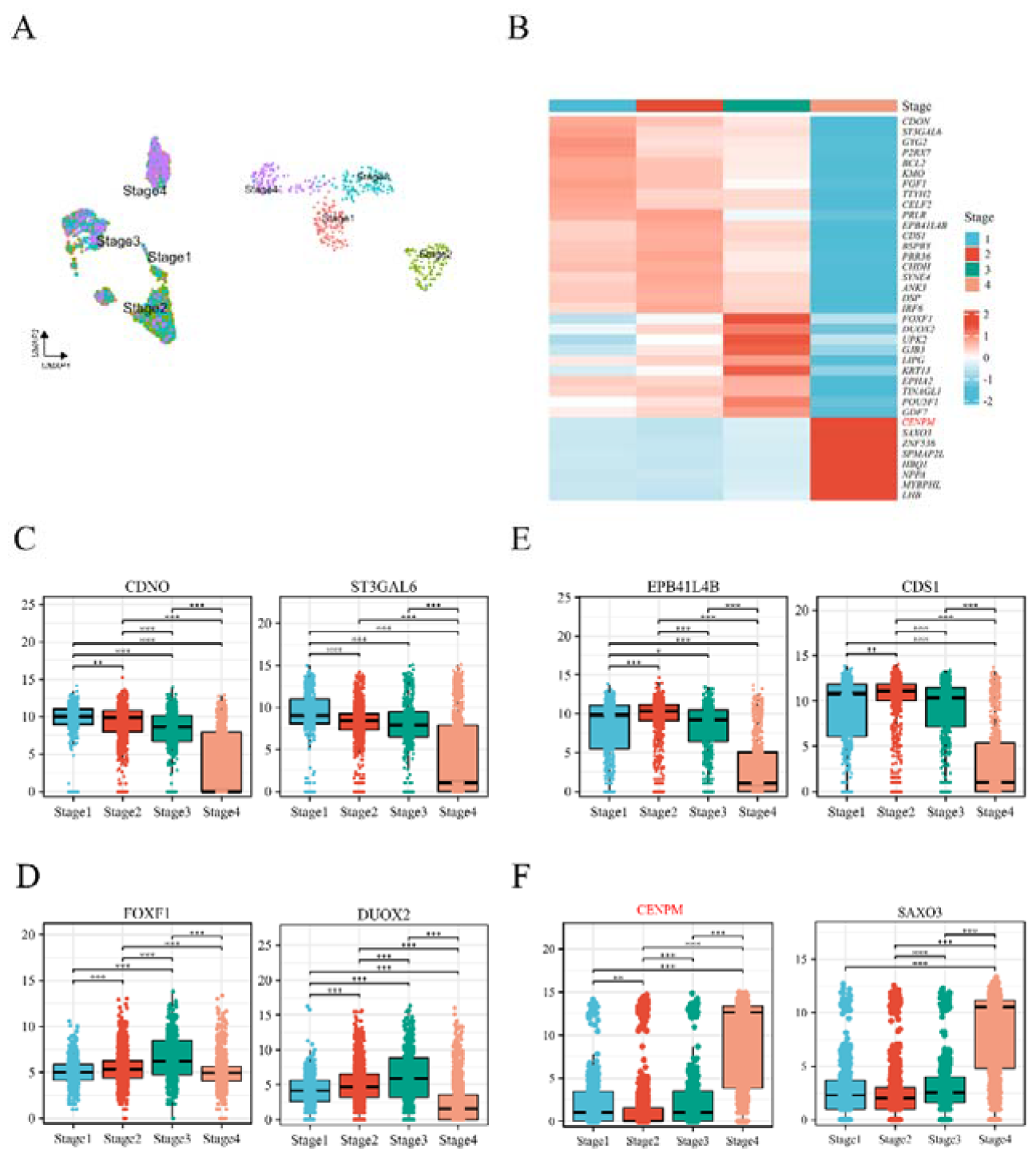
Identifying Phenotype Genes Across Diseases via clGENE. A. UMAP visualizations demonstrate clGENE’s capability in processing biological data noise, with the left side showing the pre-processed state and the right side depicting post-clGENE processing outcomes. B. Heatmaps reveal the expression patterns of genes filtered by clGENE across various lymph node metastasis phenotypes within a pan-cancer dataset. C-F. Display the expression profiles of key genes across different stages of lymph node metastasis.

### clGENE Identifying Critical Genes Bridging Similar Phenotypes Across Diverse Diseases

In our investigation, utilizing the clGENE analytical tool, we conducted an in-depth exploration of the characteristics of lymph node metastasis staging within a pan-cancer dataset. The analysis identified CENPM as a critical gene within multiple gene clusters during the fourth stage of lymph node metastasis, as depicted in (Figure 4A). CENPM, also known as Proliferation associated nuclear element 1 (PANE1), is categorized within the kinetochore protein family and was initially discovered in mouse mammary epithelial cells[12]. As a protein that connects chromosomes to microtubules, kinetochore proteins play an essential role in ensuring the accurate separation of chromosomes during cell division[13]. Furthermore, CENPM is involved not only in the binding to microtubules and the regulation of chromosome separation during cell division but also plays a role in various biological functions of the cell cycle[14, 15]. The expression of CENPM and the assembly of the kinetochore require strict regulation throughout the cell cycle, with errors in this regulation potentially leading to aneuploidy. Current research has highlighted a close association between CENPM and other genes related to tumors with similar characteristics, such as CENPA[16], CENPE[17], and CENPF. These studies further demonstrate that the overexpression of the kinetochore protein family significantly impacts the proliferation and invasion capabilities of tumors.

**Figure 4:**
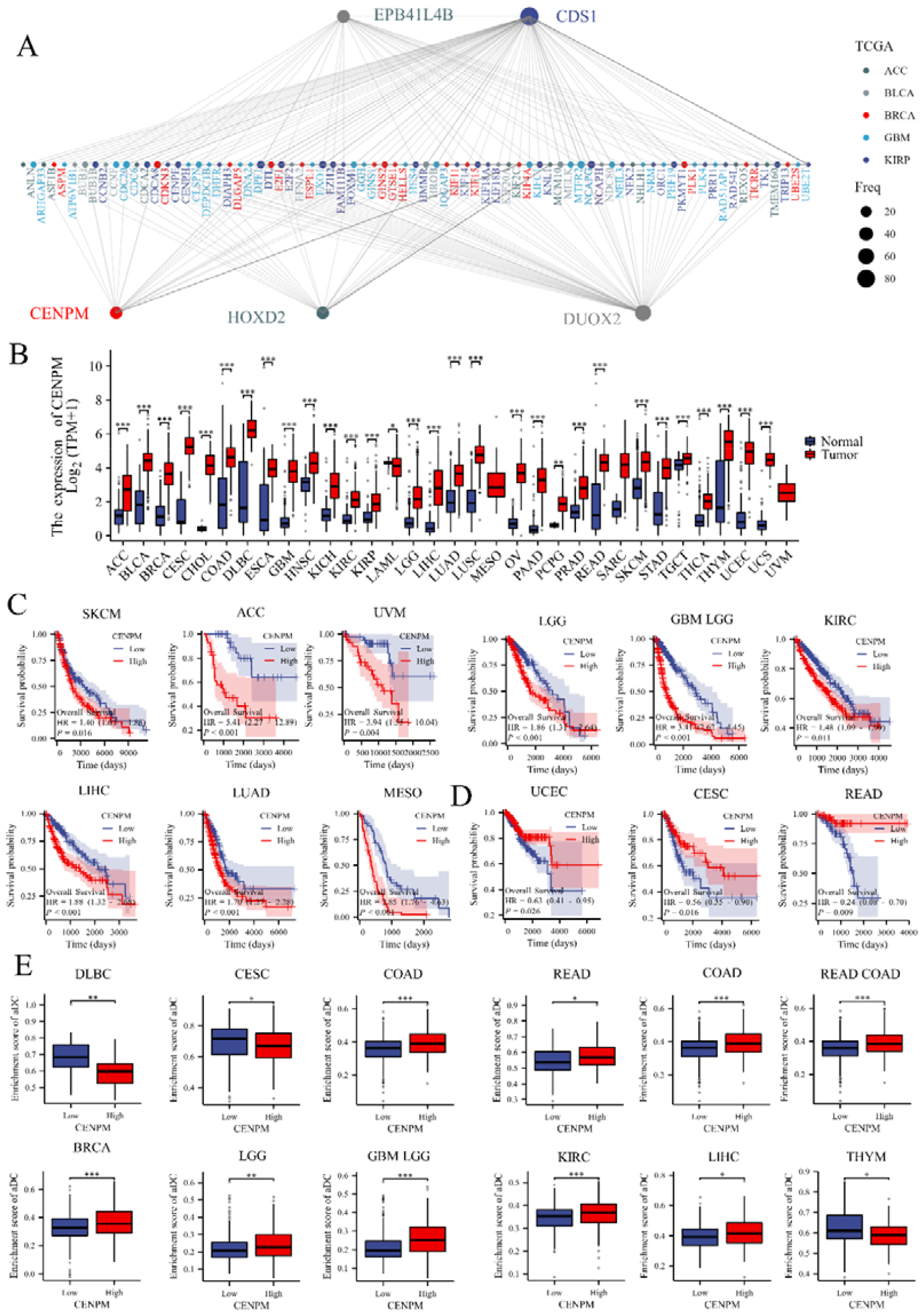
Identifying Key Genes Mediating Disease Phenotypes via clGENE Analysis. A. Through the utilization of the clGENE tool, we identified key gene clusters at various stages of lymph node metastasis across different tumor types within a pan-cancer dataset, with significant genes prominently highlighted. B. Employing this comprehensive cancer dataset, we analyzed the expression variation of CENPM across diverse types of cancer. C.D Furthermore, we assessed the impact of CENPM expression levels on the prognostic survival outcomes across various cancers. E. The comparison of immune cell infiltration scores between high and low CENPM expression groups was conducted. For all statistical analyses, a two-tailed T-test was employed, with p-values less than 0.05 considered statistically significant, and p-values less than 0.001 considered highly statistically significant.

In subsequent studies, we observed a widespread upregulation of CENPM across various cancer types (Figure 4B). Further analysis of the relationship between the expression levels of CENPM and patient survival rates revealed a significant association: elevated expression of CENPM is closely linked to poor prognosis in multiple cancers (Figure 4C). However, in specific types of cancer such as endometrial carcinoma (UCEC), cervical squamous cell carcinoma (CESC), and rectal adenocarcinoma (READ), an inverse trend was identified (Figure 4D). Moreover, in the context of immune-related analyses, the expression of CENPM was found to correlate with the immune infiltration status in various cancers (Figure 5D).

**Figure 5:**
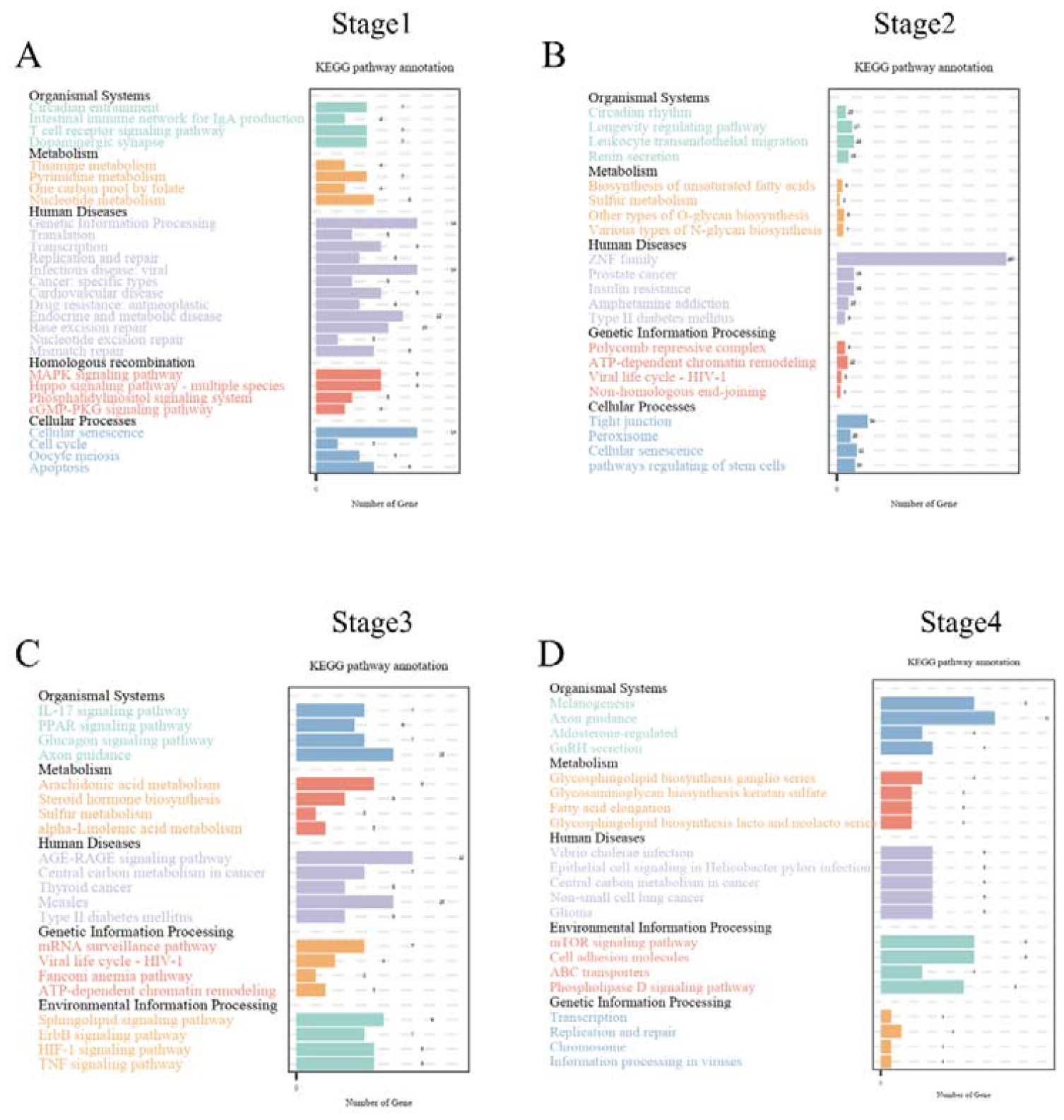
clGENE can identify key events promoting phenotypic changes in disease. A-D, KEGG enrichment analysis of key gene clusters at different stages of cancer metastasis: A for the first stage, B for the second stage, C for the third stage, and D for the fourth stage.

### clGENE can identify key events that promote phenotypic changes in disease

To explore the functional roles of gene clusters identified by clGENE, we conducted a KEGG enrichment analysis on these genes. During the early stages of lymph node metastasis (as shown in Figure 5A), we observed significant enrichment in pathways such as T cell receptor signaling, Pyrimidine metabolism, and the MAPK signaling pathway. There is ample evidence suggesting that T cell exhaustion plays a pivotal role in facilitating cancer progression[18]. Moreover, Pyrimidine metabolism is a critical pathway for the synthesis of purine and pyrimidine molecules, which are essential for DNA replication, RNA synthesis, and cellular bioenergetics[19]. The upregulation of nucleotide metabolism supports the uncontrolled proliferation of tumors, marking a hallmark of cancer. The Mitogen-Activated Protein Kinase (MAPK) signaling pathway serves as a key regulatory module in a myriad of cellular processes, including cell proliferation, differentiation, and stress response[20]. These events are all closely related to the early stages of cancer.

Interestingly, a significant discovery by clGENE in the context of lymph node metastasis was the identification of a substantial number of ZNF genes during the second phase of metastatic progression (Figure 5B). This finding aligns with the broader understanding of the epithelial to mesenchymal transition (EMT) as a pivotal mechanism in the early steps of tumor metastasis[21]. In the third phase of cancer lymphatic metastasis, we observed a significant enrichment of genes in the IL17 signaling pathway, with abundant evidence strongly suggesting a close association between the IL17 signaling pathway and cancer metastasis[22] (Figure 5C). In the study of cancer metastasis, particularly in its fourth phase, the discovery by clGENE further reinforces the understanding of the key role played by the mTOR signaling pathway in promoting cancer metastasis. This finding resonates with the previous review on the role of mTOR in cancer progression, highlighting the central role of the mTOR signaling pathway in regulating cancer cell proliferation, survival, and resistance to drug treatments (Figure 5D)[23].

### clGENE can identify key immune cells that promote similar phenotypic changes across different diseases

Changes in immune cells often predict disease progression[24]. In this study, we utilized clGENE to identify key immune cells that promote similar phenotypic changes across different diseases, further exploring their role in accelerating disease progression. Our findings reveal that, in the early stages of lymph node metastasis, active T cells, particularly CD4 and CD8 T cells, serve as the primary immune cells, aligning with previous research results. Consistently, extensive prior research has also indicated that the exhaustion of T cells marks a significant indicator of cancer exacerbation[25]. In the fourth stage of lymph node metastasis, we observed a significant presence of mast cells and memory cells as the main immune infiltrating cells. Numerous studies have already established a correlation between mast cells and cancer metastasis. In summary, our research demonstrates that clGENE can identify crucial changes in immune cells based on clinical data related to disease progression (Figure 6A,B)[25, 26].

**Figure 6:**
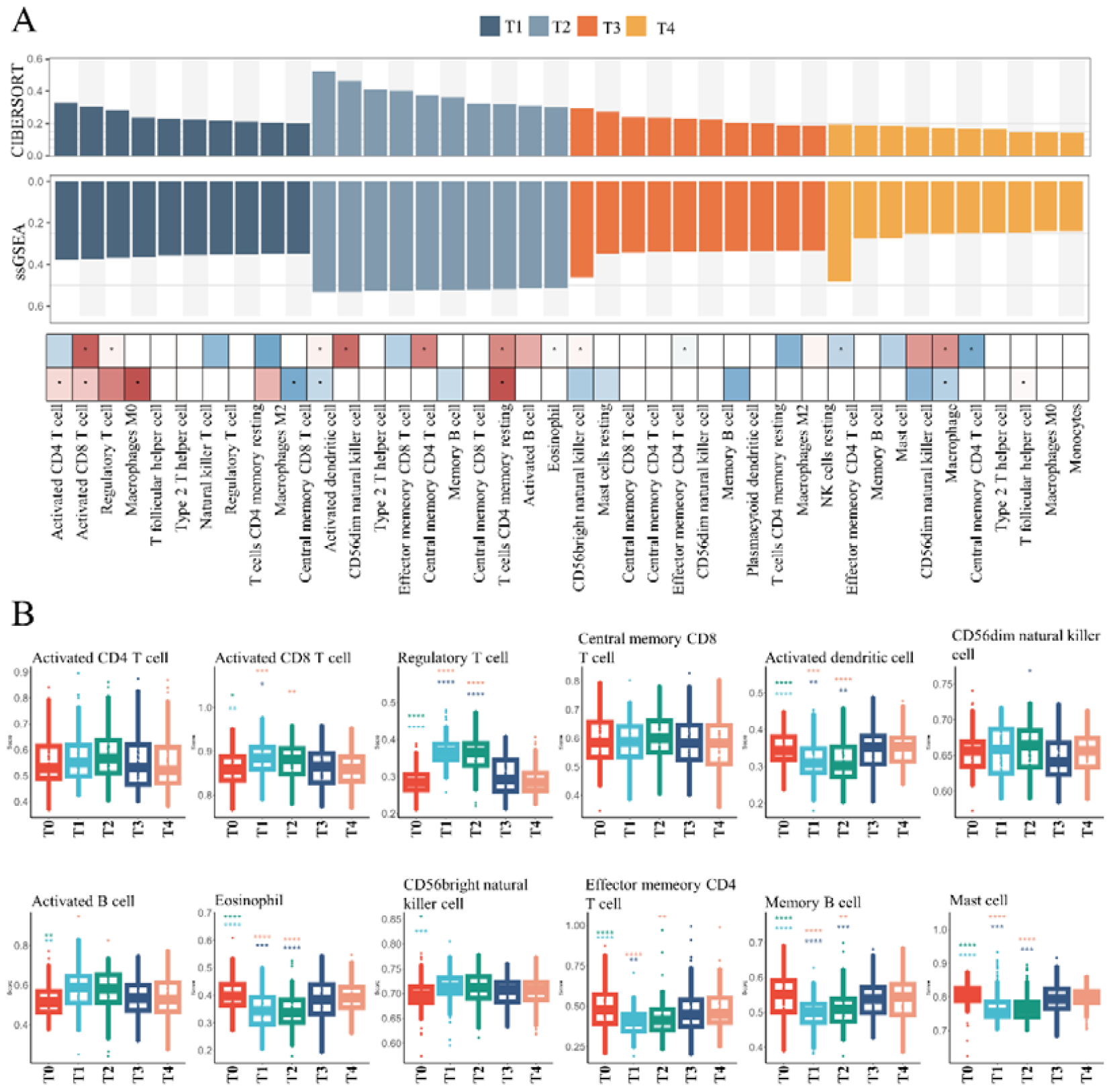
Application of clGENE to immune infiltration datasets. A utilized clGENE to analyze two immune datasets, ssGSEA and CIBERSORT, with different colors representing various stages of lymph node metastasis. The heatmap below displays immune cells that underwent significant changes. B presents the variations of key immune cells across different stages of lymph node metastasis.

## Discussion

In the combined analysis of various omics and clinical data, we discovered that clGENE is capable of precisely identifying key genes mediating similar phenotypic changes across different diseases. Furthermore, clGENE supports multifaceted data integration analysis, such as the combination of omics data with immune cell infiltration data. In the task of filtering key genes associated with similar phenotypes, clGENE delivers accurate and robust results. Additionally, clGENE efficiently integrates data from different diseases into a two-dimensional space, facilitating the batch processing of new multi-omics data. Through UMAP visualization, we observed that clGENE demonstrates exceptional performance in dimensionality reduction and batch data correction, supporting the discovery of more precise biological insights.

In recent years, although numerous multi-omics integration frameworks have been proposed, to our knowledge, there currently exists no framework specifically designed to filter key molecular mechanisms under similar phenotypes across different diseases by integrating omics with clinical data. The key advantage of the clGENE framework lies in its effective batch correction of data and precise identification of phenotype-specific expressional molecules, benefits that stem from its specially designed algorithmic components, such as deep L1 regularization and cosine similarity algorithms, inherently suited for the characteristics of omics data. Furthermore, clGENE supports parallel analysis across multiple omics, providing researchers with a powerful tool to navigate the ever-growing volume of multi-omics data, enabling the efficient extraction of critical information.

## Author contributions

Zheng Li designed the overall research strategy and wrote the manuscript. Lili Sun and Lizhi Peng manuscript was checked. All authors contributed to the article and approved the submitted version.

## Code availability

The clGENE package, implemented in R, is accessible at https://github.com/lizheng199729/clGENE, where the source code can be obtained, and the package can be downloaded for utilization.

